# The formation and content of odor memory from childhood

**DOI:** 10.64898/2026.07.16.738999

**Authors:** Jules Dejou, Moustafa Bensafi, Anne Didier, Nathalie Mandairon

**Author notes:** Equal contribution (Anne Didier, Nathalie Mandairon).

## Abstract

As illustrated by Marcel Proust’s madeleine, childhood olfactory autobiographical memory carries a stronger positive emotional valence than memories associated with other senses. However, little is known about its content or the conditions of encoding and maintenance. Here, we asked 647 participants to freely recall and characterize their earliest childhood olfactory memory. We found that this memory is typically encoded around age six, involves pleasant odors embedded within rich multisensory contexts, and retains a strong positive emotional valence despite age-related declines in some experiential qualities. The most frequently reported scenes took place at home, in the presence of the mother, and were predominantly associated with food-related odors, although natural-vegetation odors were also commonly reported. By contrast, the non-olfactory memory is encoded earlier, visually dominated, less multisensory, less pleasant, and more frequently associated with institutional settings and friends. These findings characterize childhood olfactory memory and highlight its distinctive content among autobiographical memories.

## Introduction

Autobiographical memory is defined as the recollection of personal events drawn from one’s own life combining episodic elements (specific events in time and space and the relationships between them) and semantic elements (general knowledge about oneself and the world) (Conway, 2005). Among autobiographical memories, it is well known that olfactory memories from childhood are capable of bringing back an entire scene from one’s childhood that one thought had long been forgotten. This particular form of memory, in which the olfactory sense plays a central role, was famously described by Marcel Proust in his work *In Search of Lost Time* (1913). Since then, the subjective sensations associated with childhood olfactory memory have been studied using various methodologies, primarily drawing on the cueing-method (Galton, 1879; Rubin, 1982). This approach involves presenting words, images, odors or multimodal cues to participants, who are then asked to recall a memory evoked by these cues and to evaluate it across several dimensions. Other approaches involve studying these sensory memories without presenting any stimuli, either through mental imagery (Schlintl et al., 2022; Willander & Larsson, 2008) or by requesting regular reports (the diary method) regarding significant or early memories (Bogenschütz et al., 2022; Hutmacher, 2021).

Using these various techniques, olfactory memory has been described following the acronym LOVER, for “Limbic”, “Old”, “Vivid”, “Emotional” and “Rare” (Larsson et al., 2014, but see Hackländer et al., 2019 for a recent meta-analysis) compared to memories related to other sensory modalities. As regards the age of childhood olfactory memories, the ‘peak of reminiscence’ was reported to occur at an earlier stage (i.e., before the age of 10) than that of non-olfactory memories (Chu & Downes, 2000; Cornell Kärnekull et al., 2020; Goddard et al., 2005; Miles & Berntsen, 2011; Willander et al., 2015; Willander & Larsson, 2006, 2007, 2008). The second hallmark of childhood olfactory memory is its strong positive emotional valence compared with non-olfactory memory (Arshamian et al., 2013; Chu & Downes, 2002; de Bruijn & Bender, 2018; Herz, 2004; Herz & Schooler, 2002; Miles & Berntsen, 2011; Rubin et al., 1984; Willander & Larsson, 2007). However, the literature shows some inconsistencies in the subjective feelings evoked by the olfactory memory, as not all the studies replicated all or part of the LOVER acronym. Most of these studies use the cueing-method, with olfactory stimulation, a methodology that may bias participants’ reports and introduce confounding effects related to sensory stimulation itself (Hackländer et al., 2019; Lopis et al., 2023). Some other studies used spontaneous evocation of memories, without sensory cues, and did not show a greater emotional load for odor-associated memories compared to memories associated with the other senses (Bogenschütz et al., 2022; Hutmacher, 2021). Finally, we previously reported using spontaneous recall that childhood olfactory memory typically arises from repeated experiences (more than 5 times for 73% of subjects; Dejou et al., 2026).

While some facets of olfactory memory have been extensively studied (cue efficiency, age of the memory, emotional load, vividness, feeling of being brought back in time), we feel that there is room to improve our knowledge of the content and circumstances in which this memory is formed (i.e., which odor, when, where, with whom) and the subjective feelings it evokes when recalled. Thus, our objective was to investigate the spontaneous recall of the earliest childhood olfactory memory without olfactory cues, thereby avoiding biases linked to sensory stimulation and ensuring retrieval of memories most relevant in terms of age and personal significance. Using a large-scale online survey, 647 participants were asked to recall both their earliest olfactory and non-olfactory memory. They provided a written description (including location, people present, associated perceptions and emotions) and answered several questions (1–9 scales, single choice, or open-ended) to characterize the sensory content of the memory, the encoding circumstances, the maintenance of the memory over time, and the emotional impact of its recall. This approach allowed identifying the ‘recipe’ for creating a childhood olfactory memory, assessing its emotional significance in adulthood and highlighting its specific features in comparison with non-olfactory memory.

We found that the earliest childhood olfactory memory is typically encoded around six years of age and is associated with pleasant odors experienced in rich multisensory contexts. The most represented autobiographical scenes took place at home and in the presence of the mother. Across these memories, the remembered odors were predominantly food-related, followed by odors from nature and vegetation. These memories retain a strong positive emotional valence throughout life despite age-related declines in some experiential qualities. By contrast, the earliest non-olfactory memory is encoded earlier, is visually dominated, less multisensory, less family-centered, and overall less pleasant.

## Results

### Characteristics and emotional content of childhood olfactory memory

Participants first reported their age at the time of the remembered event. The age distribution of the earliest olfactory memory was bimodal, with a main peak at 5.89 years and a secondary peak at 9.95 years (mean = 6.47 years; **Fig. 1A**). To assess the emotional valence associated with this olfactory memory, participants rated the six basic emotions described by Paul Ekman (1992): happiness, sadness, fear, anger, surprise, and disgust, on a 1–9 scale. Recollection of the event was characterized by ratings significantly higher for happiness than for the negative emotions (F(3.39, 2192.74) = 845.29, p < 0.0001; **Fig. 1B; Supplementary Table 1**). Participants also rated the overall pleasantness of the memory, and, consistent with the emotional ratings, most evaluated it as highly pleasant (χ^2^(2, N = 647) = 671.52, p < 0.0001; **Fig. 1C; Supplementary Table 1**). Together, these findings indicate that the earliest childhood olfactory memory is recalled as a strongly positive emotional experience. We next verified that olfaction was indeed the predominant sensory modality in the event participants had described. To this end, participants rated the contribution of each sensory modality on a 1–9 scale, allowing us to assess the multisensory content of the memory (F(3.58, 2312.49) = 406.20, p < 0.0001; **Fig. 1D; Supplementary Table 2**). As expected, olfaction emerged as the dominant sensory component of these memories, although the recalled experiences were clearly multisensory. We then sought to characterize the odor associated with the childhood olfactory memory. Participants rated its hedonic value on a 1–9 scale, revealing that the odor was perceived as highly pleasant (χ^2^(2, N = 647) = 492.48, p < 0.0001; **Fig. 1E; Supplementary Table 1**). Notably, the hedonic rating of the odor was strongly correlated with the overall pleasantness of the associated memory (S = 186.10^5^, p < 0.0001, r = 0.59; **Fig. 1F**). This relationship remained robust after subsampling (for each combination of memory and odor pleasantness ratings, up to five observations were randomly selected) and after excluding the high-density cluster of responses at ratings of 8:8 and 9:9 (**Supplementary Fig. 1**).

**Figure 1.**
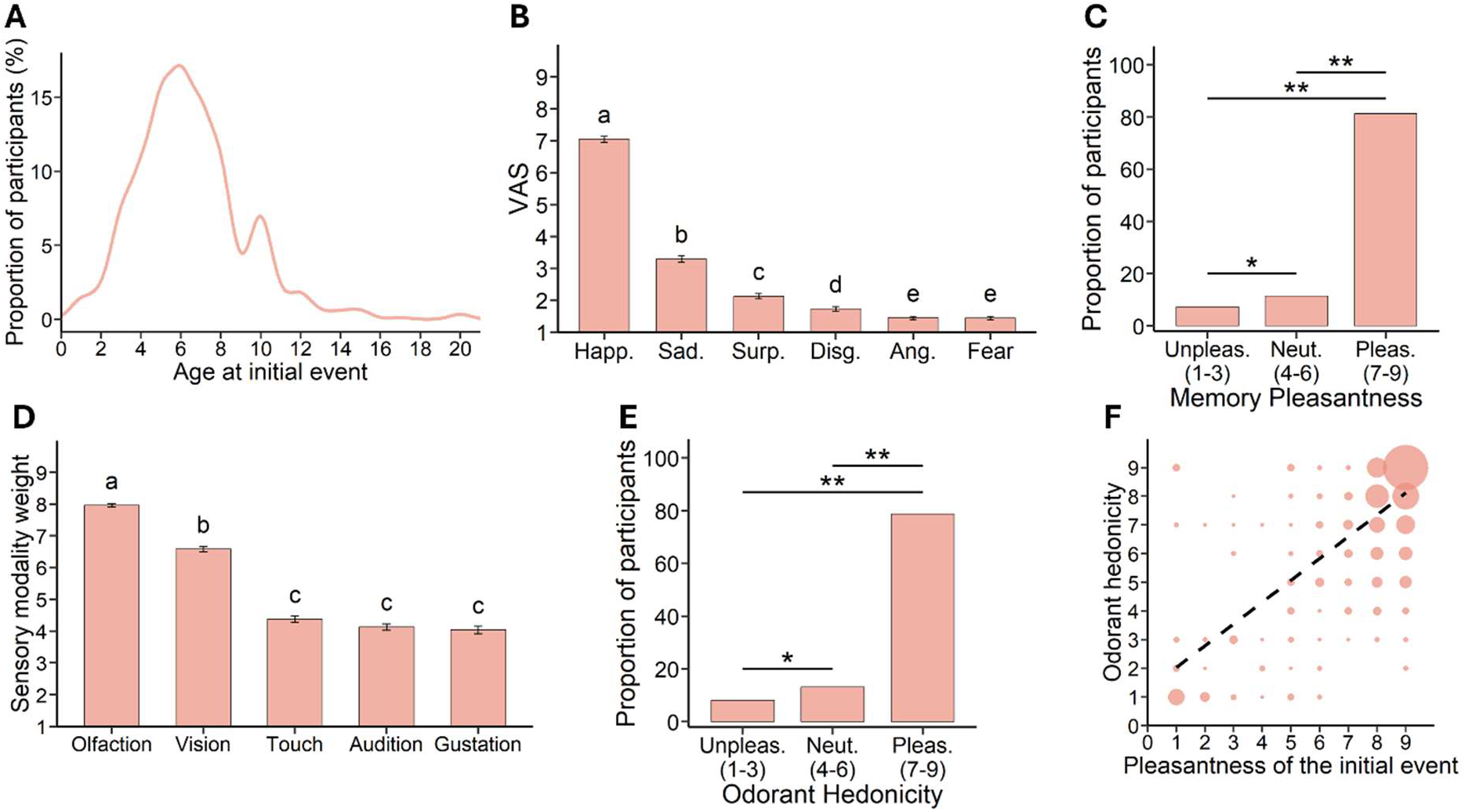
Characteristics of childhood olfactory memory. **(A)** Age at occurrence of the childhood olfactory memory, showing a bimodal distribution with peaks at 5.89 and 9.95 years. **(B)** Emotional profile of the childhood olfactory memory at recall, characterized by predominant happiness compared with negative emotions. **(C)** Distribution of childhood olfactory memories according to their pleasantness, showing a predominance of happy memory. **(D)** Relative contribution of each sensory modality to the childhood olfactory memory, showing the predominance of olfaction within a rich multisensory experience. **(E)** Distribution of childhood odors according to their hedonic value, showing a predominance of pleasant odors. **(F)** Positive correlation between the hedonic value of the odor and the pleasantness of the associated childhood memory. Data are represented as individual data points and mean ± SEM. Statistical significance is indicated as *p < 0.05, **p < 0.01. Letters indicate statistical groups identified by ANOVAs and post hoc multiple-comparison tests; significance was set at p < 0.05.

To assess the specificities of olfactory memory, we compared earliest childhood olfactory memory with the earliest non-olfactory memory, which participants were also asked to recall. The age distribution of non-olfactory memory peaked at 4.02 years (mean = 5.85 years; **Fig. 2A, left**), significantly earlier than that of olfactory memory (t(646) = −5.01, p < 0.0001) (**Fig. 2A, right**). These results indicate that the earliest olfactory memory is not necessarily the oldest autobiographical memory. In addition, we observed that the age of earliest olfactory memory was independent of participants’ current age (S = 422.10^5^, p = 0.11, *r* = 0.064), whereas the age of earliest non-olfactory memory increased with participants’ age (S = 386.10^5^, p = 0.0002, *r* = 0.15) (**Supplementary Fig. 2**). This suggests that olfactory memory is more stably anchored within an individual’s life history. The emotions associated with non-olfactory memory were predominantly happiness, followed by surprise and negative emotions (F(2.77, 1787.85) = 357.98, p < 0.0001; **Supplementary Table 1**). However, compared with olfactory memory, non-olfactory memory was characterized by lower ratings of positive emotions and higher ratings of negative emotions (one-sample t-tests on the log-ratio, FDR-corrected, F(2.29, 1481.31) = 75.115, p < 0.0001; **Fig. 2B; Supplementary Table 1**). We also found that vision was the dominant sensory modality in non-olfactory memory (F(3.58, 2314.12) = 582.25, p < 0.0001; **Fig. 2C; Supplementary Table 2**), in contrast to the olfactory dominance observed in olfactory memory (two-way repeated-measures ANOVA, interaction between memory type and sensory modality, F(3.74, 2413.07) = 473.19, p < 0.0001; **Supplementary Table 2**). Interestingly, non-olfactory memory contained weaker contextual sensory details, as indicated by lower cumulative sensory modality ratings compared with olfactory memory (t(646) = −14.9, p < 0.0001; **Fig. 2D**). Non-olfactory memory also differed from olfactory memory in the frequency with which the original event was repeated, with a greater proportion of participants reporting a unique event (p < 0.0001) and a smaller proportion reporting repeated occurrences of similar events (p < 0.0001) compared with childhood olfactory memory (Dejou et al., 2026; same set of participants) (**Fig. 2E; Supplementary Table 3**). We then examined the subjective experience elicited during recall by asking participants to rate their feelings of nostalgia, mental time travel (the feeling of being brought back in time), and memory vividness on 1–9 scales. Compared with non-olfactory memory, childhood olfactory memory was associated with significantly stronger feelings of nostalgia (t(646) = −10.1, p < 0.0001; **Fig. 2F**), a greater sense of being transported back in time (t(646) = −6.19, p < 0.0001; **Fig. 2G**), and higher vividness, although the latter effect was more modest (t(646) = −3.46, p = 0.0006; **Fig. 2H**).

**Figure 2.**
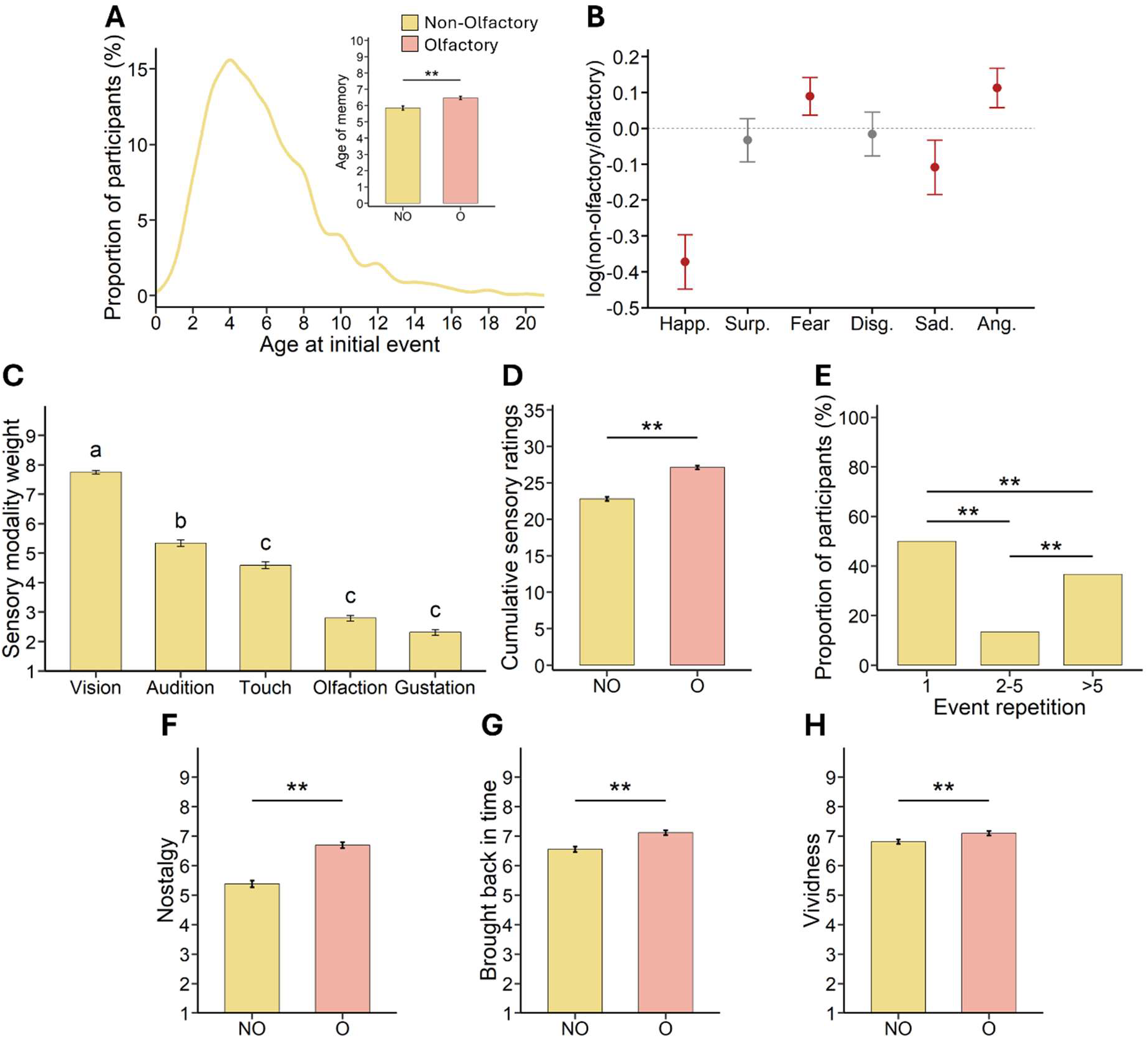
Distinctive features of childhood olfactory memory relative to non-olfactory memory. **(A)** Age at occurrence of the earliest childhood non-olfactory memory, showing a distribution peaking at 4.02 years. **(B)** Emotional profile of the childhood non-olfactory memory at recall compared to that of olfactory memory (log(ratio non-olfactory/olfactory)). **(C)** Relative contribution of each sensory modality to the childhood non-olfactory memory, showing the predominance of vision. **(D)** Cumulative sensory ratings, indicating richer multisensory content in childhood olfactory memory than in non-olfactory memory. **(E)** Frequency of repetition of the initial event associated with childhood non-olfactory memory, showing that about half of the participants reported a unique event. **(F)** Childhood olfactory memory is associated with greater nostalgia, **(G)** a stronger feeling of being brought back in time, and **(H)** higher vividness than non-olfactory memory. Data are represented as individual data points and mean ± SEM or proportions, as appropriate. Statistical significance is indicated as *p < 0.05, **p < 0.01. Letters indicate statistical groups identified by ANOVAs, post hoc multiple-comparison tests or proportion comparison tests; significance set at p < 0.05.

To summarize, the earliest childhood olfactory memory typically arises from repeated experiences (>5 repetitions; Dejou et al., 2026) occurring around the peak age of 6 years. It is associated with both a pleasant odor and a pleasant context, and its recall predominantly evokes happiness. In contrast, the earliest non-olfactory memory is typically formed around the peak age of 4 years, requires fewer repetitions, and is recalled less positively than olfactory memory. Moreover, compared with non-olfactory memory, olfactory memory is more nostalgic, produces a stronger feeling of mental time travel, and is experienced as more vivid.

### What do we remember?

We then sought to determine whether certain smells or categories of smells are more commonly associated with olfactory memories than others. We thus analyzed the list of odors reported by the participants to the question “What is the main smell associated with this memory?”. As the olfactory perceptive space is highly multidimensional and there is no universal odor classification, we used artificial intelligence (AI) (see Methods) prompted to categorize the odors with no other specific instruction. Eight categories were identified based on the odor sources. One describing wood odors was added to the category Nature and Vegetation, leading to 7 categories presented below in the order of importance (χ^2^(6, N = 613) = 292.94, p < 0.0001; **Fig. 3A; Supplementary Table 4**). Olfactory memory was predominantly composed of food-related odors, among which bread, sugar, toast, coffee, milk and chocolate are the most frequently cited (**Fig. 3B**). The second most represented category comprised nature and vegetation odors including grass, flowers, pine and hay (**Fig. 3C**). Other, less frequent categories include home, industry-related odors, perfumes and personal care products, animal and human body odors, and rain odors. Next, we proceeded with the analysis of the participants’ written recollections of their olfactory memory. Using AI (see Methods), we first constructed a dictionary containing the words (nouns, adjectives and verbs) of the corpus. Then, AI built eight domains in a semi-supervised manner, in which the words were classified (see Methods, **Supplementary Fig. 3**). We then conducted a detailed analysis of the domains sorted by AI “Social”, “Places” and “Subjective feelings” corresponding to the instructions given to the participants in the questionnaires (where were you, with whom and what were your feelings) (**Fig. 3D-F**). Regarding the “Social” domain, we revealed that the mother is the most cited person, followed far behind by the father and the brother, and even further behind by other family members or friends (**Fig. 3D, Supplementary Table 5**). When analyzing the “Places” where the memorized event took place, we found that words referring to the home are the most represented, followed, in descending order, by natural environments, vacation places, school and hospital (**Fig. 3E, Supplementary Table 6**). Olfactory memory was also characterized by a strongly positive emotional valence, dominated by words describing feelings of joy and well-being (**Fig. 3F; Supplementary Table 7**). Thus, rather than being randomly distributed across childhood experiences, the earliest olfactory memory consistently converged toward a few selective social, contextual, and emotional features. While olfactory memory was more strongly centered on the nuclear family, non-olfactory memory more frequently involved friends (**Fig. 3G, Supplementary Table 5**). Non-olfactory memory was also more often embedded in institutional settings, particularly hospital or school (**Fig. 3H; Supplementary Table 6**), and was associated with a higher proportion of negative feelings, such as surprise, fear and sadness (**Fig. 3I; Supplementary Table 7**). Finally, we examined the semantic organization of childhood memories by constructing binary co-occurrence networks from participants’ written recollections of the childhood olfactory and non-olfactory memories they had recalled. Pearson correlation matrices were computed between the domains “Place”, “Social” and “Food” (**Fig. 3J-K**). The childhood olfactory memory network formed a single, highly interconnected component linking food, social figures and place. Within this network, Mother, Home and Meals emerged as highly connected nodes linking multiple categories. In contrast, the childhood non-olfactory memory network displayed a more modular organization, comprising two main clusters: one centered on food-related categories (Meals, Eating/drinking, Sweet, and Fruits) and another centered on social figures and school-related settings, with School and Teachers acting as the main organizing nodes.

**Figure 3.**
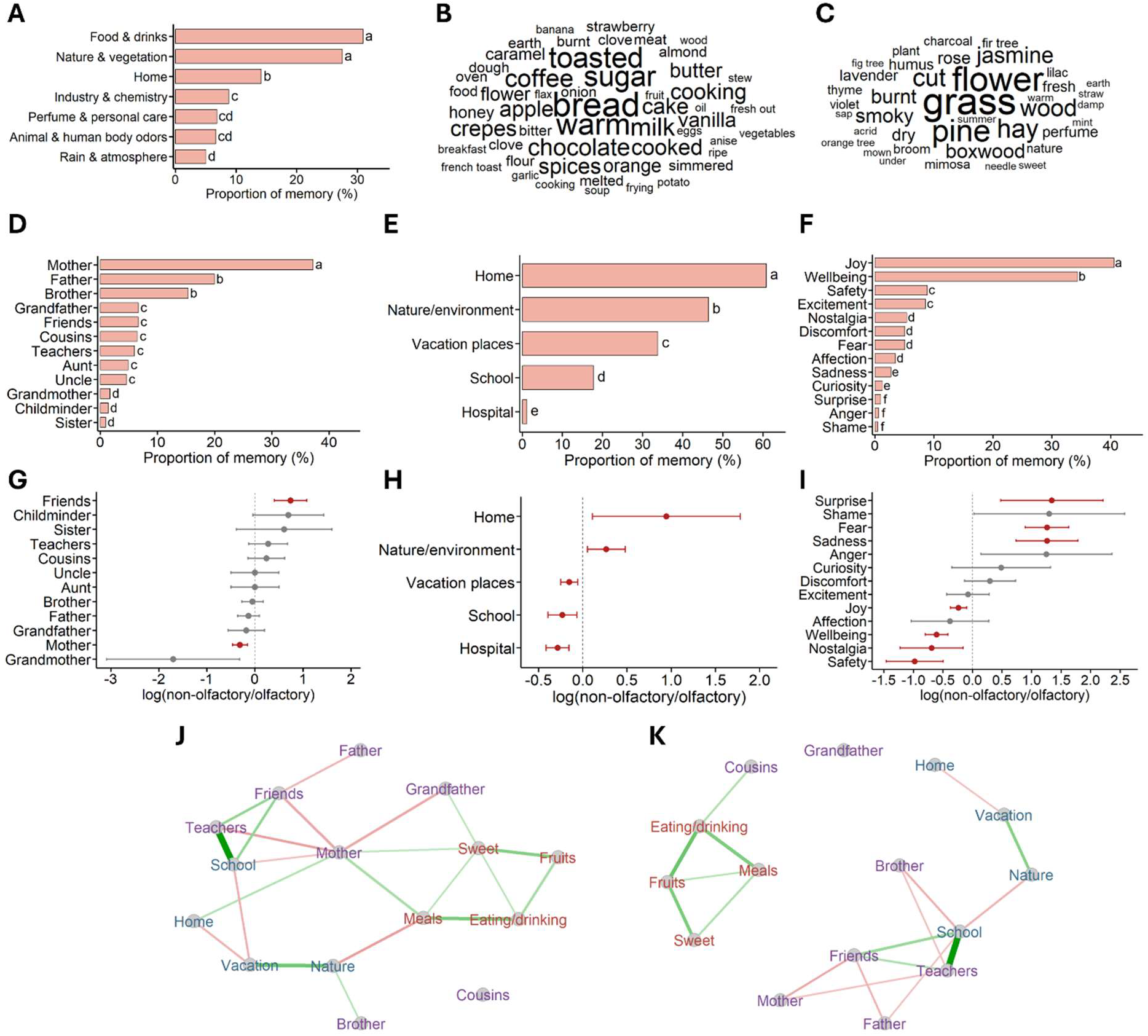
Content of childhood olfactory memory. **(A)** Distribution of the main odor categories associated with childhood olfactory memory, showing a predominance of food and drink odors followed by nature and vegetation odors. **(B)** Word cloud of odors from the Food & drinks category, with bread, sugar, toast, coffee, milk and chocolate being the most frequently reported. **(C)** Word cloud of odors from the Nature & vegetation category, with grass, flowers, pine and hay being the most frequently reported. **(D)** The distribution of social context categories reveals a predominance of the mother figure in childhood olfactory memory. **(E)** Distribution of the places in which childhood olfactory memories were formed, with home being the most frequent one. **(F)** Distribution of the emotions associated with childhood olfactory memory, showing a predominance of joy and well-being. **(G–I)** Forest plots comparing the relative representation of **(G)** social figures, **(H)** places, and **(I)** emotions in childhood olfactory versus non-olfactory memory (log ratio non-olfactory/olfactory). Values < 1 indicate greater representation in olfactory memory, whereas values > 1 indicate greater representation in non-olfactory memory. **(J–K)** Semantic networks generated from participants’ written recollections of childhood olfactory **(J)** and non-olfactory **(K)** memories. Nodes represent odor categories, social figures, settings and contextual descriptors, whereas edges indicate their co-occurrence within the same memory. The childhood olfactory memory network forms a highly interconnected structure linking food categories, social figures, and places. In contrast, the childhood non-olfactory memory network is more fragmented, with one cluster centered on food and another centered on social figures and school-related settings. Different letters indicate significant differences between categories following proportion comparison tests.

Taken together, these findings reveal the detailed content of the earliest childhood olfactory memory: it is preferentially formed during emotionally positive, food-related experiences occurring at home in the presence of primary attachment figures, particularly the mother. This combination is unique to olfactory memory.

### Childhood olfactory memory throughout life

We have previously shown that the childhood odor associated with the earliest olfactory memory is re-encountered throughout life at frequencies ranging from never to once a week (Dejou et al., 2026). We therefore investigated to what extent re-exposure to this odor spontaneously evokes the associated autobiographical memory. We found a markedly non-uniform distribution of memory recall frequency among participants (χ^2^(3, N = 530) = 240.04, p < 0.0001; **Fig. 4A**). Notably, the largest proportion of participants (42.2%) reported that exposure to their childhood odor “always” evoked the associated memory. Fewer participants reported that the odor “often” or “sometimes” evoked the memory, whereas only a small proportion (4%) indicated that it “never” triggered memory recall (**Fig. 4A; Supplementary Table 8**). Interestingly, the ability of the odor to evoke the childhood memory depended on how frequently it had been encountered throughout life. Odors encountered infrequently (“yearly” or “less than yearly”) were the most likely to evoke the associated childhood memory. In contrast, odors encountered more frequently (“monthly” or “weekly”) were significantly less effective at triggering memory recall (**Fig. 4B; Supplementary Table 8**). Together, these findings indicate that repeated lifetime exposure to a childhood odor progressively reduces its ability to evoke the associated autobiographical memory.

**Figure 4.**
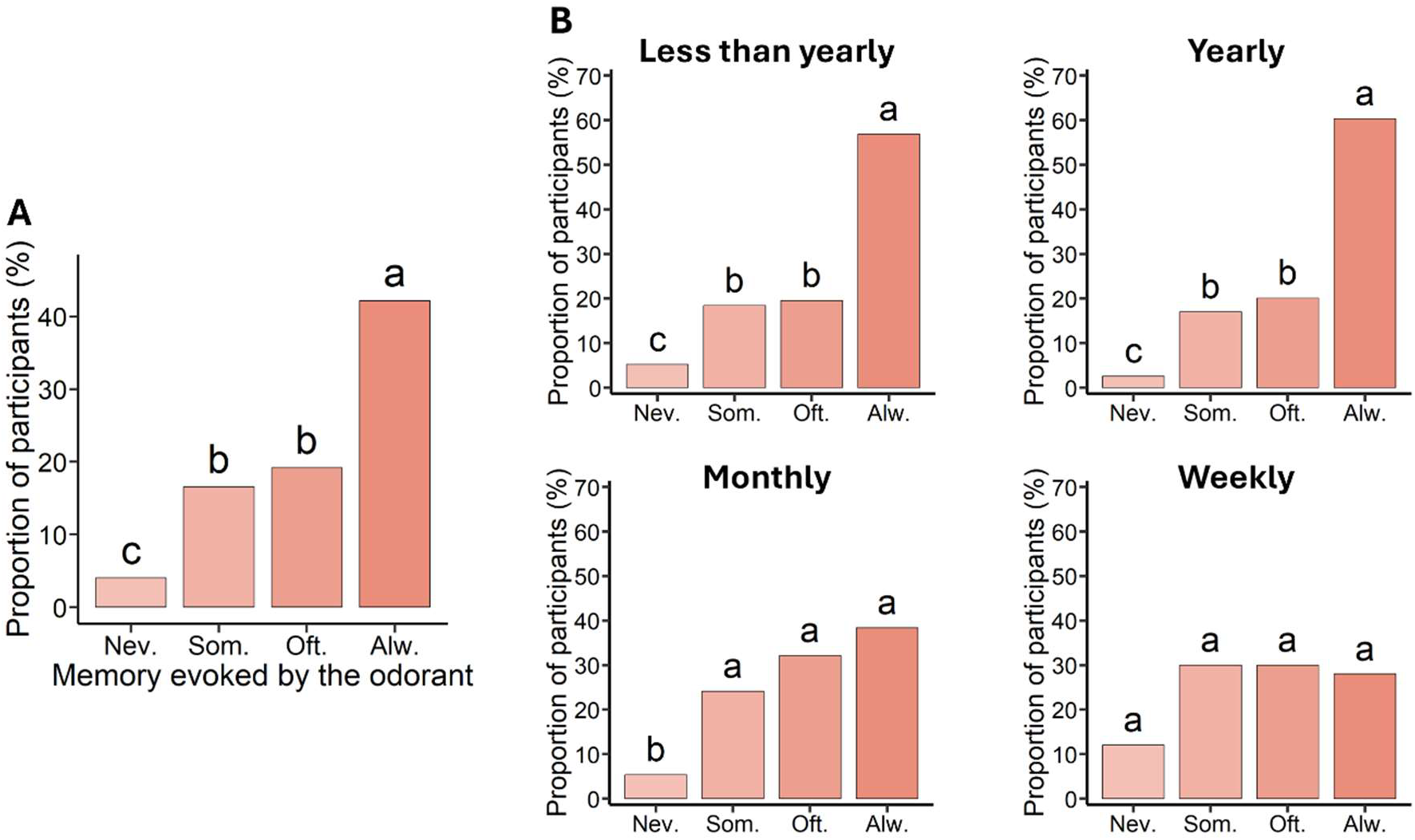
Lifetime odor exposure shapes odor-evoked memory recall. **(A)** Distribution of participants according to the frequency with which re-exposure to their childhood odor evoked the associated autobiographical memory. **(B)** Memory-evoking ability of the childhood odor according to its lifetime exposure frequency. Rarely encountered odors (less than yearly or yearly) were more likely to always evoke the associated autobiographical memory than odors encountered monthly or weekly. Data are represented as proportions. Different letters indicate significant differences between categories following proportion comparison tests, significance was set at p < 0.05.

### The subjective properties of the childhood olfactory memory are affected by the age at recall

We examined the effects of the age of participants on the subjective characteristics of the childhood olfactory memory by dividing participants into three age groups: 18-40, 41-60 and 61-90. To compare throughout time the overall emotional valence of the memory, we used a composite emotional score calculated by averaging positive emotion ratings and reverse-scored negative emotion ratings at the time of recall. We found that the overall emotional valence of the olfactory memory declined with age (F(2, 644) = 4.155, p = 0.016; post hoc comparison: 41–60 vs. 61–90, p = 0.004; **Fig. 5A**), indicating a less positive emotional evaluation of the memory in older adults. To determine which specific emotions accounted for this age-related decline, we then grouped participants into two age categories (18–60 and 61–90 years), as the composite emotional score did not differ between the 18–40 and 41–60 groups (p = 0.226). A mixed two-way ANOVA revealed an age-dependent change in the emotional profile of olfactory memories (emotion × age interaction: F(3.39, 2186.5) = 9.78, p < 0.0001; **Supplementary Fig. 4**). Post hoc analyses showed that the decline in the composite emotional score with aging was driven exclusively by an increase in negative emotion ratings, whereas positive emotion ratings remained unchanged (**Supplementary Fig. 4, Supplementary Table G**).

**Figure 5.**
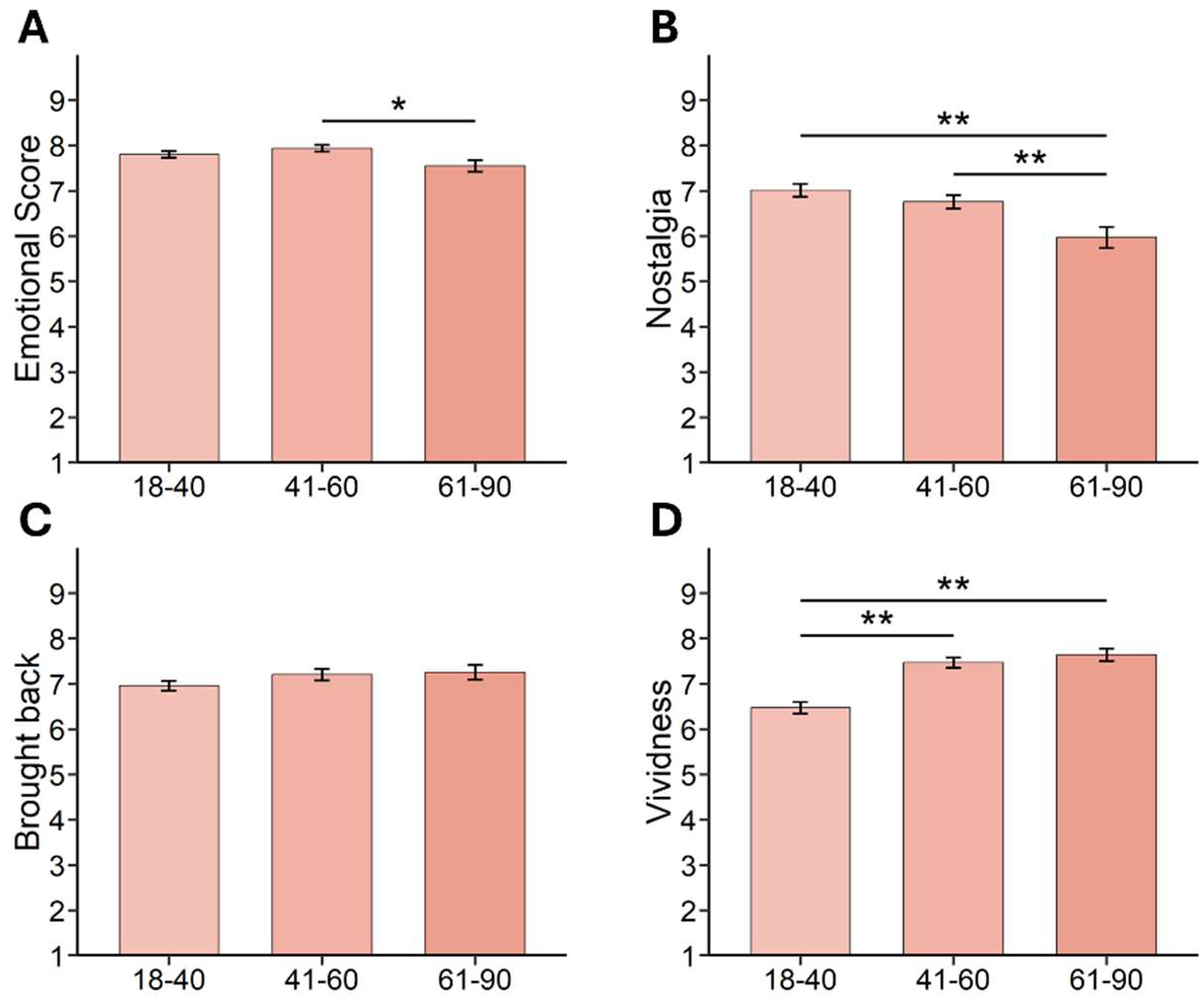
The subjective experience of childhood olfactory memory depends on participants’ age. Increasing age is associated with lower **(A)** composite emotional valence score and **(B)** feeling of nostalgia but not **(C)** feeling of being brought back in time. **(D)** Conversely, the feeling of vividness increases with age (18-40, n=266; 41-60, n=238; 61-90, n=143). Data are represented as data points and mean ± SEM. Statistical significance depicted as *p < 0.05, **p < 0.01.

In addition, feelings of nostalgia declined with age (F(2,644) = 8.823, p = 0.0002; post hoc comparison: 41–60 vs. 61–90, p = 0.006; **Fig. 5B**). In contrast, participants’ age had no effect on the feeling of being brought back in time (F(2,644) = 1.551, p = 0.213; **Fig. 5C**). Interestingly, the vividness of childhood olfactory memories increased with age (F(2,644) = 25.022, p < 0.0001; **Fig. 5D**). Specifically, vividness ratings were higher in the 41–60 group than in the 18–40 group (p < 0.0001). To determine whether these age-related effects were specific to childhood olfactory memory, we performed the same analyses on the earliest non-olfactory memory. In contrast to olfactory memories, participants’ age had no significant effect on either the composite emotional valence (F(2,644) = 1.298, p = 0.27) score or nostalgia (F(2,644) = 1.201, p = 0.30). However, both the feeling of being brought back in time (F(2,644) = 4.20, p = 0.015; post hoc comparison: 18-40 vs. 41-60, p = 0.016, 18-40 vs. 61-90, p = 0.015, 41–60 vs. 61–90, p = 0.73) and memory vividness (F(2,644) = 10.53, p < 0.0001; post hoc comparison: 18-40 vs. 41-60, p = 0.016, 18-40 vs. 61-90, p < 0.0001, 41–60 vs. 61–90, p = 0.136) increased with age (**Supplementary Fig. 5**).

### Determinants of the positive valence of childhood olfactory memory

Next, we investigated whether the key characteristics of childhood olfactory memory (i.e., high olfactory and gustatory weights, rich multisensory content, and repetition of the initial event) contributed to its positive emotional valence. We found that greater olfactory weight (S = 377.10^5^, p < 0.0001, r = 0.16; **Fig. 6A**), richer sensory content (i.e., the sum of sensory modality ratings; S = 378.10^5^, p < 0.0001, r = 0.16; **Fig. 6B**), and repeated exposure to the initial context (F(2,644) = 11.876, p < 0.0001; post hoc comparisons: 1 vs. 2-5, p = 0.128, 1 vs. >5, p < 0.0001, 2-5 vs. >5, p = 0.035; **Fig. 6C**) were all associated with greater pleasantness of the childhood olfactory memory.

**Figure 6.**
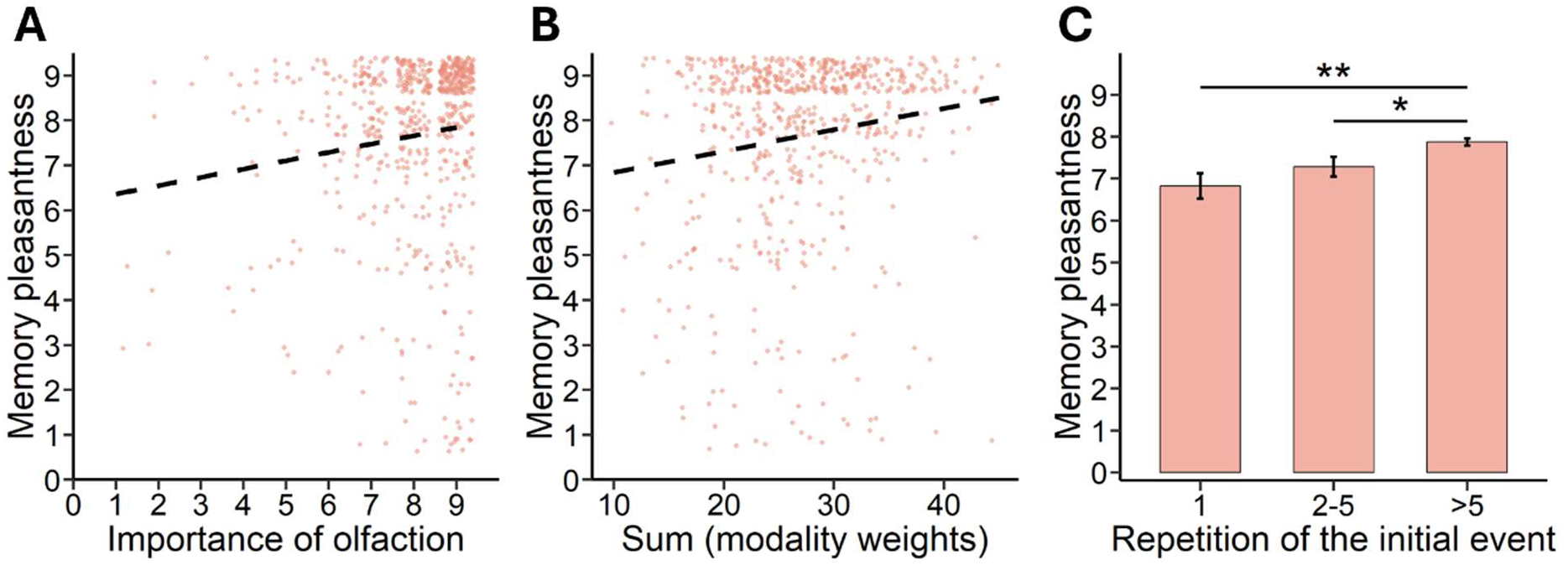
The singular features of childhood olfactory memory directly affect its emotional valence. The emotional valence of the childhood olfactory memory is correlated with **(A)** the olfaction weight and **(B)** the cumulative sensory ratings. It is also positively impacted by **(C)** the number of repetitions of the initial event. Data are represented as data points and mean ± SEM. Statistical significance depicted as *p < 0.05, **p < 0.01.

To determine whether the effect of olfactory content on the pleasantness of childhood olfactory memory was specific to olfaction itself, rather than simply reflecting a richer sensory experience, we performed a mediation analysis (to test whether the relationship between a predictor and an outcome is direct or mediated by a third variable). We found that the contribution of olfaction to memory pleasantness was both indirect, through its association with greater sensory richness (p = 0.0004), and direct (p = 0.035), indicating that olfaction enhances the emotional valence of the memory through a specific mechanism independent of overall sensory content (**Supplementary Table 10**). In contrast, analyses of the other sensory modalities (vision, audition, gustation, and touch) revealed only an indirect effect on memory pleasantness via increased sensory richness. For the non-olfactory memory, gustation directly and positively contributed to memory pleasantness (p < 0.0001), with a comparable trend for olfaction (p = 0.075), whereas vision, the sensory modality that dominates the description of the memory, did not positively contribute to the pleasantness of memory (**Supplementary Table 10**). Thus, chemosensory senses – olfaction and gustation – positively influence the emotional valence of both olfactory and non-olfactory remembered experiences.

## Discussion

While the properties of childhood olfactory memory have been widely reviewed in the literature, little was known about the content of the memory and the circumstances that allow this memory to be formed and maintained in the long term. We found that childhood olfactory memory is made of a pleasant odor related to food or nature, experienced several times around the age of 6, often with the mother, and triggers happiness that does not fade with time unless the odor is frequently re-encountered throughout life.

Regarding the age at which the memory was formed, we found it to be later for the olfactory memory compared to the non-olfactory one (6 vs. 4 years), which is in line with Hutmacher and colleagues (2021) but contradicts some literature dating olfactory memory earlier than visual memory (Chu & Downes, 2000; Cornell Kärnekull et al., 2020; Goddard et al., 2005; Miles & Berntsen, 2011; Willander & Larsson, 2006). This could be explained first by a biased estimation of the age of the earliest visual memory which depends on the question asked to the subjects (Jack & Hayne, 2007). Indeed, spontaneous, non-cued recollection methods allow the remembering of earlier memory than those oriented by sensory cues (Jack & Hayne, 2007; Willander & Larsson, 2006). Second, the older age of olfactory memory compared to non-olfactory one may be explained by the different circumstances and contents of the memories. Most olfactory memory results from a repetition (at least 5 times, Dejou et al., 2026) of the memorized event, suggesting a routine spanning over a longer period of time compared to the unique event forming the majority of non-olfactory memories. Our study further brings new information related to the content of olfactory childhood memory. Based on the reports by the participants of breakfast or meals odors, the home environment and the presence of the mother, one may propose that the cultural food habits in France, from which our participants originated, may influence memory formation in children reaching the age when they actually sit at the table of these meals. Besides cultural aspects, our results are in line with Schlintl and colleagues (2022) who reported that most of the olfactory memories dating from childhood were pleasant. Furthermore, these three elements (food, home, mother) may provide reward and feelings of safety making the experience highly positive and allowing its memorization as an adaptive mechanism. Altogether, these findings strongly suggest that most childhood olfactory memories are emotionally positive and further reveal that this could arise from the specific content of olfactory memory. However, it would be interesting to further assess the cultural contribution to the contents of early olfactory memory. Finally, although not necessarily older, the olfactory memory seems to be enduring and resistant to the passage of time, as the age of participants did not influence the dating of the memory, unlike non-olfactory memory.

The lifelong persistence of this memory may stem from the fact that the memory’s dominant odor is encoded as a unified representation within a broader sensory landscape, rather than as a set of decomposable subcomponents (Hackländer et al., 2019; Yeshurun & Sobel, 2010). As previously mentioned, our findings also revealed that the olfactory memory is associated with greater sensory information in comparison to the non-olfactory memory, which is in line with previous studies (Chu & Downes, 2002; Toffolo et al., 2012). This was partly due to an increased gustatory component, which confirms the strong proximity between these two sensory modalities, notably in defining the food flavor. This also suggests that some of our olfactory memories are in fact olfacto-gustatory, although their proportion remains to be determined. Importantly, the amount of sensory information affects the emotional valence of the memory: the more sensory detail the memory contains, the more pleasant it is. Yet olfaction alone influenced valence independently of this multisensory effect. The hypothesis of an integrated olfactory landscape embedded in olfactory memory may also explain why common odors such as toasted bread still evoke emotional old memories: while they are frequently re-encountered during life, they may not occur with other odors of the initial context which could counteract the attenuation with re-exposure of the evocative power of odors.

Regarding the level of initial event repetition, this parameter has rarely been studied, yet the available evidence indicates that olfactory memory more often originates from repeated events than non-olfactory memory (**Fig. 2E;** Dejou et al., 2026; Ernst et al., 2021; Goddard et al., 2005). Here, we reproduced and extended these results showing that this parameter strongly influences the emotional properties of our memory. Interestingly, the episodic memory has classically been referred to as the memory of unique events, and not repeated ones (Tulving, 1972, 1983). Thus, sensory-related autobiographical memory is commonly investigated by asking participants to recall memories of unique events, as for episodic memory, which then may lead to the recall of less pertinent olfactory memory. Recently, some authors have proposed the existence of a continuum between episodic and semantic memory, along which the repeated life events would belong to an intermediate category called “personal semantic memories” (Addis & Szpunar, 2024; Bontkes et al., 2025; Renoult et al., 2012, 2019). In the future, it appears important to assess the impact of event repetition on the characterization of autobiographical memory rather than controlling it, especially in the context of olfactory memories.

All these singular encoding properties (age of formation, sensory richness, content specificity, repetition of initial event) confer unique emotional properties to our childhood olfactory memory. Indeed, we confirmed that this memory is associated with a highly positive valence, as evidenced by higher ratings for positive emotions and lower ratings for negative ones, as well as a stronger feeling of nostalgia, of being brought back in time and of vividness compared to the earliest non-olfactory memory, corroborating previous findings (Hackländer et al., 2019; Larsson et al., 2014). Importantly, our approach did not rely on olfactory stimulation to evoke the memory, and it also enabled fine-grained correlation analyses of how these factors contribute to the emotional valence of the olfactory memory.

Interestingly, our findings reveal that the positive emotional load of childhood olfactory memory, as well as its nostalgia, decreases with increasing participants’ age. These results are surprising in light of a well-documented effect, named the age-related positivity effect, that is characterized by a tendency of older adults to recall more positive olfactory and non-olfactory autobiographical memories (Comblain et al., 2005; Reed et al., 2014; Ros & Latorre, 2010; Schlagman et al., 2006, 2009; Yamamoto & Sugiyama, 2023). This difference could be related to the non-cued method used here which may require more complex search strategies that were reported to suppress the age-related positivity effect (Reed et al., 2014; Schlagman et al., 2009; Siedlecki et al., 2015).

During childhood, one experiences thousands of odors, raising the question of what makes an odor special and remembered for a lifetime. By clarifying the content and the conditions under which childhood olfactory memory is most likely to be formed and preserved as well as how it is retrieved in adulthood, this work, using large-scale free-recall methodology, contributes to answering this question by a better understanding of the distinctiveness of childhood olfactory memory. This distinctiveness relies on the memory of an odor within a sensory landscape and a specific social and spatial context.

## Material G Method

### Participants

The participants were recruited through various channels including research volunteers mailing lists, CNRS and local weekly newspapers. All participants provided informed consent. The study was approved by the Ethical Review Committee of Inserm (no. 23-1002, April 4, 2023). All collected data were anonymized. In total, 652 participants were recruited to complete an online questionnaire. Five participants reported a history of an olfactory disorder during childhood and were excluded from the analysis. Thus, the final sample consisted of 647 participants (174 men and 473 women), aged 18 to 85 years (M = 45.4, SD = 17.2).

### Procedure

The online questionnaire was hosted in LimeSurvey. The participants were instructed to recall and describe their earliest olfactory memory, followed by their earliest non-olfactory memory. In the first section, participants were instructed to recall an odor that marked their childhood, going back as far as possible. After retrieving an olfactory autobiographical memory, they were asked to provide a written description of it using 200-800 characters specifying the location, the people present, and the associated perceptions and emotions. Participants then had to rate this memory on several self-reported measurements (single choice, 1-9 Likert scales or open-ended), grouped into three categories: the event that was memorized, the recollection process and the associated odor. In the second section, participants were asked to recall another childhood memory, this time without strong olfactory component, again going back as far as possible. It followed the same structure as the first section, except for the questions related to the odor, which were omitted. A detailed list of all assessed measures can be found in **Table 1**.

**Table 1.**
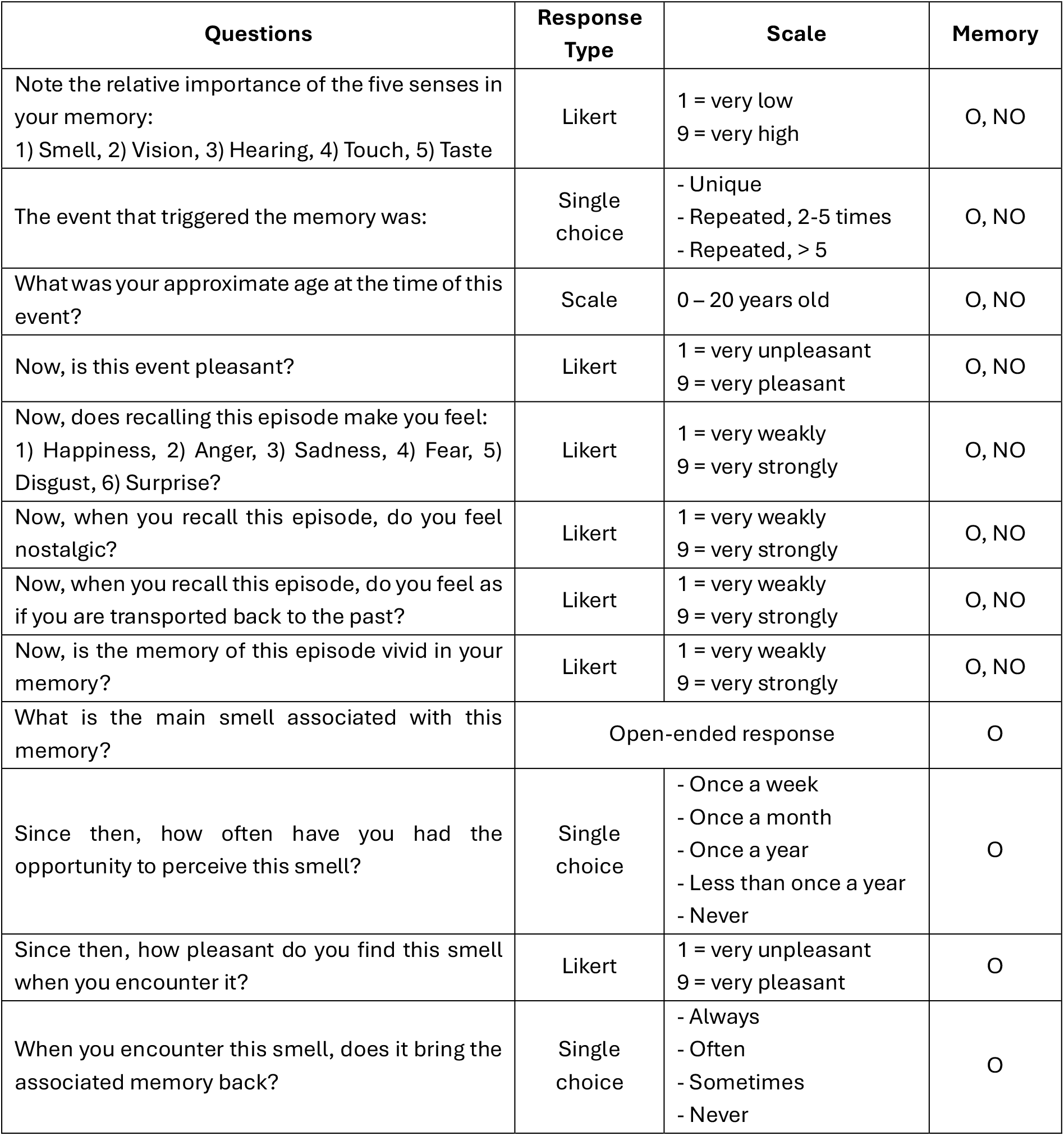
List of assessed variables. O = Olfactory memory; NO = Non-Olfactory memory

| Questions | Response Type | Scale | Memory |
| --- | --- | --- | --- |
| Note the relative importance of the five senses in your memory:<br>1) Smell, 2) Vision, 3) Hearing, 4) Touch, 5) Taste | Likert | 1 = very low<br>9 = very high | O, NO |
| The event that triggered the memory was: | Single choice | - Unique<br>- Repeated, 2-5 times<br>- Repeated, > 5 | O, NO |
| What was your approximate age at the time of this event? | Scale | 0 – 20 years old | O, NO |
| Now, is this event pleasant? | Likert | 1 = very unpleasant<br>9 = very pleasant | O, NO |
| Now, does recalling this episode make you feel:<br>1) Happiness, 2) Anger, 3) Sadness, 4) Fear, 5) Disgust, 6) Surprise? | Likert | 1 = very weakly<br>9 = very strongly | O, NO |
| Now, when you recall this episode, do you feel nostalgic? | Likert | 1 = very weakly<br>9 = very strongly | O, NO |
| Now, when you recall this episode, do you feel as if you are transported back to the past? | Likert | 1 = very weakly<br>9 = very strongly | O, NO |
| Now, is the memory of this episode vivid in your memory? | Likert | 1 = very weakly<br>9 = very strongly | O, NO |
| What is the main smell associated with this memory? | Open-ended response |  | O |
| Since then, how often have you had the opportunity to perceive this smell? | Single choice | - Once a week<br>- Once a month<br>- Once a year<br>- Less than once a year<br>- Never | O |
| Since then, how pleasant do you find this smell when you encounter it? | Likert | 1 = very unpleasant<br>9 = very pleasant | O |
| When you encounter this smell, does it bring the associated memory back? | Single choice | - Always<br>- Often<br>- Sometimes<br>- Never | O |

### Odor classification

Each participant freely provided a short label describing the odor associated with their earliest childhood olfactory memory. Using a large language model (Claude Opus 4.7), we derived a data-driven categorization scheme that yielded eight categories, which we reviewed and refined into seven categories. Specifically, a category describing wood odors was merged into the category Nature & vegetation. The 7 final categories were: *Food & drinks, Nature & vegetation, Home, Industry & chemistry, Perfumes & personal care, Animal & human body odors, Rain & atmosphere.* Each label was automatically assigned to a single category. Entries that were truncated, illegible, or too vague to interpret (e.g., “impossible to describe”; *n* = 34 out of 647) were labeled *Unclassified* and excluded from category-level analyses. For each corpus, we reported the proportion of odor labels falling into each category. For example, a value of 30.9% for *Food & drinks* indicates that 30.9% of the named odors (and therefore participants) were classified as food- or drink-related. Because each label was assigned to a single category, these proportions were mutually exclusive and summed to 100%.

### Word clouds

To illustrate the lexical content of the two most frequent odor categories, we generated word clouds for the *Food & drinks* and *Nature & vegetation* categories. For each category, the odor labels provided by all participants were concatenated into a single corpus and tokenized into individual words. Stopwords were removed, and morphological variants of the same term (e.g., “apple” and “apples”) were manually merged into a single form. The frequency of each remaining term was then counted, and the resulting frequencies were displayed as a word cloud in which font size is proportional to the number of occurrences.

### Text classification

To compare content of olfactory and non-olfactory memory, we characterized the participants’ free-recall narratives using a lexicon-based content-analysis approach. A large language model (ChatGPT 5.5) generated an initial set of 6 domains relevant to the description of autobiographical memory (Family/close persons, Domestic places, Nature/outdoors, Food, Body/sensations, Objects), which we then refined into eight final domains: *Sensoriality, Place, Social, Subjective feelings, Time, Food, Body,* and *Object*. Within each domain, a set of more specific categories was defined to allow finer-grained classification. We then built a lexicon mapping words to these domains and categories. Through several iterative rounds of human-model collaboration, candidate words proposed by the model were reviewed, corrected, expanded, and validated, yielding a comprehensive lexicon in which each entry was assigned to one category and its associated domain. Prior to classification, both corpora (olfactory and non-olfactory memories) were preprocessed: spelling errors were automatically corrected and each narrative was tokenized into individual words. Each token was then compared against the dictionary. When a token matched a dictionary entry, the participants’ narrative was labeled with the corresponding category and domain. Overall, the dictionary matched 18.4% of tokens in the olfactory corpus and 15.9% of tokens in the non-olfactory corpus. In contrast to the odor classification, this classification operated at the token level and was non-exclusive: a single narrative could be assigned to several domains and categories, and each domain was counted only once per participant regardless of the number of matching tokens. For each corpus, we computed the prevalence of every domain and category as the proportion of participants whose narrative contained at least one word matching that domain or category. Because a given narrative could match several domains, these proportions were not mutually exclusive and did not sum to 100%.

### Olfactory category co-occurrence networks

We constructed association networks for the three domains *Place, Social* and *Food*, separately for the olfactory and non-olfactory corpora. Within each corpus, categories with extreme prevalence (below 5% or above 95%) were excluded, as they carry little associative information. Pairwise associations were quantified using the phi coefficient (φ) (i.e., the Pearson correlation between binary variables). To retain only substantial associations, edges with |φ| < 0.10 were set to zero. To allow a direct comparison between memory types, the two networks were then restricted to the set of categories present in both corpora. Networks were visualized using a force-directed layout, with nodes colored by domain.

### Data analysis

Data analyses were performed using R (version 4.5.3). As the participants described their olfactory and non-olfactory autobiographical memory, one-tailed or two-tailed paired t-tests or Wilcoxon tests, with Welch correction when needed, were performed to compare olfactory and non-olfactory memory. Additionally, one-way ANOVAs were used to assess the effect of participants’ age on subjective memory properties, mixed two-way ANOVAs to analyze the effect of participants’ age on emotion ratings, and repeated-measures ANOVAs to examine the emotion ratings and the relative importance of sensory modalities across memory types. *Post-hoc* comparisons were conducted when needed, with Bonferroni corrections applied. Proportion tests (for the comparison of qualitative variables), Spearman correlations, and mediation analyses (to examine whether the effect of an independent variable on a dependent variable is transmitted through a third variable, called the mediator) were also performed. To analyze the emotional content of the memories, composite emotional valence scores were calculated as follows: [VAS(Happiness) + (10 − VAS(Sadness)) + (10 − VAS(Disgust)) + (10 − VAS(Anger)) + (10 − VAS(Fear))] / 5. Surprise was excluded from the composite score given its ambivalent valence, as it can be experienced as either positive or negative depending on context. For the corpora analysis, the prevalence of categories between olfactory and non-olfactory memory were compared using McNemar’s test with continuity correction. As a measure of effect size, we computed the ratio of the two prevalences (olfactory relative to non-olfactory). Because ratios are asymmetric on a linear scale, confidence intervals were derived on the log scale using the delta method, with a covariance term accounting for the within-participant correlation between the two memory types, and then back-transformed. The inverse ratio (non-olfactory relative to olfactory) was also reported to allow effects to be read in either direction. To account for multiple comparisons across subcategories, McNemar p-values were adjusted using the Benjamini–Hochberg false discovery rate (FDR) procedure, with adjusted p < 0.05 considered significant. A parallel ratio-based approach was used to compare emotion ratings between memory types. For every participant and emotion, we computed the ratio of the non-olfactory to the olfactory rating and log-transformed it. Participants for whom this ratio was undefined (e.g., a zero denominator) were excluded for that emotion. For each emotion, we then computed the geometric mean ratio across participants (the back-transformed mean of the individual log-ratios), together with its 95% confidence interval derived from the standard error on the log scale. Because the log-ratio equals the within-participant difference between the log-transformed ratings, its departure from zero (i.e., a ratio departing from 1, indicating no difference between memory types) was tested using a one-sample t-test, equivalent to a paired t-test on the log scale. P-values were adjusted across emotions using the Benjamini–Hochberg false discovery rate (FDR) procedure, with adjusted p < 0.05 considered significant. A log-ratio greater than zero indicates that the emotion was rated more strongly for the non-olfactory memory, and conversely.

## Acknowledgments

The author(s) declare that financial support was received for the research, authorship, and/or publication of this article. This work was supported by Observatoire B2V des Mémoires (doctoral fellowship to J.D.), CNRS, ANR-23-CZE17-0065-01, INSERM, Claude Bernard Lyon 1 University, IUF. The manuscript was edited for language and clarity with the assistance of ChatGPT.

## Author Contributions

J.D., M.B., A.D. and N.M. designed research; J.D. analyzed data; J.D., A.D. and N.M. wrote the paper.

