## Supplementary Fig. 1 for "The formation and content of odor memory from childhood"

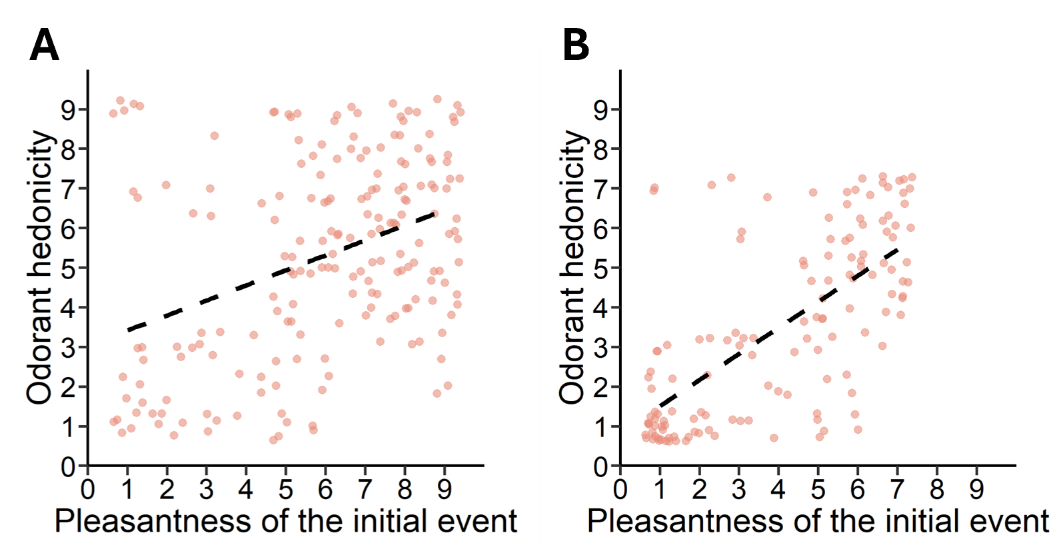
**Supplementary Figure 1.** **The pleasantness of the odorant and of the initial event are positively correlated.** The positive correlation observed between the initial event and its associated odorant is conserved following **(A)** a subsampling or **(B)** a removal of the high-cluster at the 8:8 and 9:9 intersections.
