## Supplementary Fig. 2 for "The formation and content of odor memory from childhood"

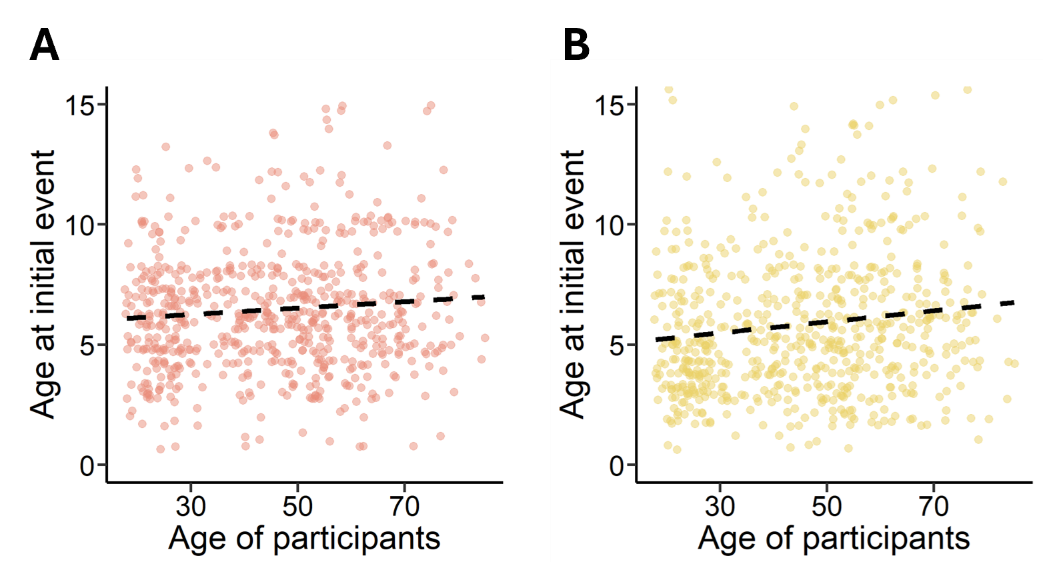
**Supplementary Figure 2. The olfactory memory is resistant to the passage of time.** The dating of the initial event depending on the age of the participants shows **(A)** no significant correlation for the olfactory memory, but **(B)** a positive correlation for the non-olfactory memory.
