## Supplementary Fig. 3 for "The formation and content of odor memory from childhood"

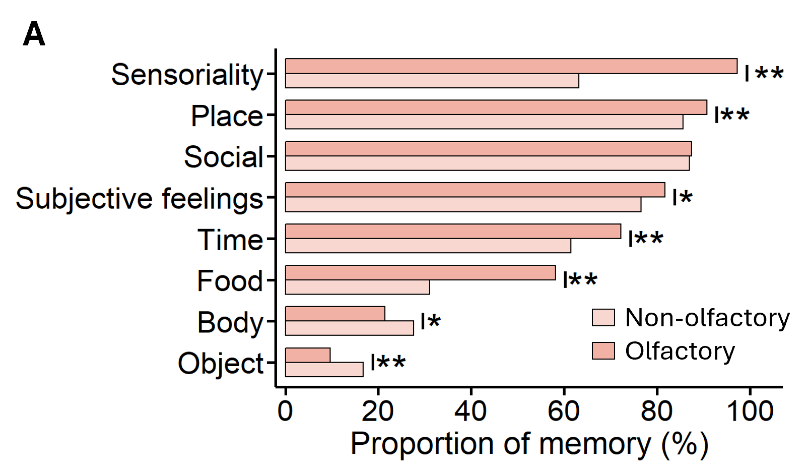
**Supplementary Figure 3. Olfactory and non-olfactory memory are built in distinct contexts. (A)** Compared with non-olfactory memory, olfactory memory shows stronger representation of sensoriality (p < 0.0001), place (p = 0.005), subjective feelings (p = 0.027), time (p < 0.0001) and food (p < 0.0001), and weaker representation of body (p = 0.013) and object (p = 0.0003). Statistical significance is indicated as *p < 0.05, **p < 0.01.
