## Supplementary Fig. 4 for "The formation and content of odor memory from childhood"

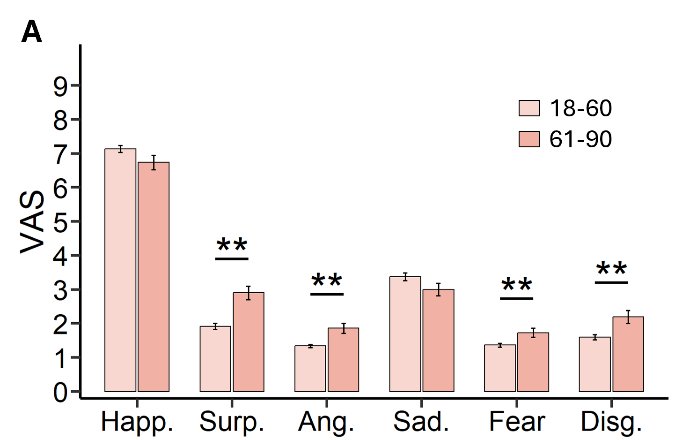
**Supplementary Figure 4. Olfactory memory becomes associated with stronger negative emotions with age. (A)** Compared with the 18-60 group (n=504), the 61-90 group (n=143) reports higher ratings for surprise (p < 0.0001), anger (p < 0.0001), fear (p = 0.0049) and disgust (p = 0.0006). Data are represented as data points and mean ± SEM. Statistical significance depicted as *p < 0.05, **p < 0.01.
