## Supplementary Fig. 5 for "The formation and content of odor memory from childhood"

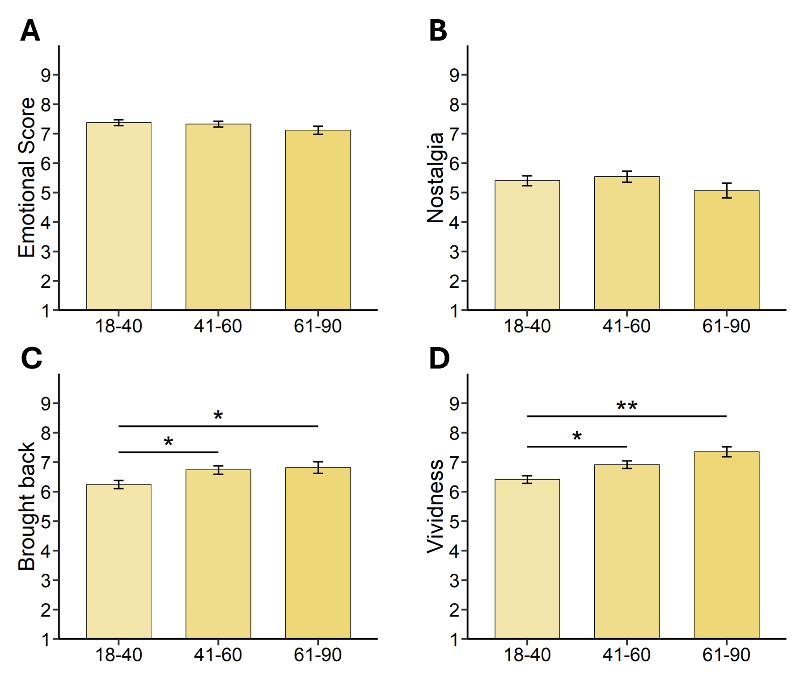
**Supplementary Figure 5. Subjective experience of childhood non-olfactory memory depending on participants’ age**. Increasing participants’ age does not influence **(A)** the composite emotional valence score nor **(B)** the feeling of nostalgia. However, it increases the feelings of **(C)** being brought back in time and **(D)** vividness (18-40, n=266; 41-60, n=238; 61-90, n=143). Data are represented as data points and mean ± SEM. Statistical significance depicted as *p < 0.05, **p < 0.01.
