## Supplementary Table 1 for "The formation and content of odor memory from childhood"

| **Olfactory memory - Emotion** | | | | | **Olfactory memory - Pleasantness** | | | |
| --- | --- | --- | --- | --- | --- | --- | --- | --- |
| **group1** | **group2** | **stat** | **df** | **p.adj** | **Comparison** | **N1** | **N2** | **p.adj** |
| Anger | Disgust | -4.23 | 646 | **0.0004** | 1-3 vs 4-6 | 47 | 74 | **0.0177** |
| Anger | Fear | 0.15 | 646 | 1 | 1-3 vs 7-9 | 47 | 526 | **< 0.0001** |
| Anger | Happiness | -47.05 | 646 | **< 0.0001** | 4-6 vs 7-9 | 74 | 526 | **< 0.0001** |
| Anger | Sadness | -18.87 | 646 | **< 0.0001** | **Olfactory memory - Odor Hedonics** | | | |
| Anger | Surprise | -7.88 | 646 | **< 0.0001** | **Comparison** | **N1** | **N2** | **p.adj** |
| Disgust | Fear | 3.97 | 646 | **0.001** | 1-3 vs 4-6 | 43 | 70 | **0.0141** |
| Disgust | Happiness | -38.03 | 646 | **< 0.0001** | 1-3 vs 7-9 | 43 | 417 | **< 0.0001** |
| Disgust | Sadness | -13.74 | 646 | **< 0.0001** | 4-6 vs 7-9 | 70 | 417 | **< 0.0001** |
| Disgust | Surprise | -3.95 | 646 | **0.001** |  |  |  |  |
| Fear | Happiness | -48.84 | 646 | **< 0.0001** |  |  |  |  |
| Fear | Sadness | -18.03 | 646 | **< 0.0001** |  |  |  |  |
| Fear | Surprise | -7.94 | 646 | **< 0.0001** |  |  |  |  |
| Happiness | Sadness | 26.42 | 646 | **< 0.0001** |  |  |  |  |
| Happiness | Surprise | 41.46 | 646 | **< 0.0001** |  |  |  |  |
| Sadness | Surprise | 9.28 | 646 | **< 0.0001** |  |  |  |  |
| **Non-olfactory vs olfactory memory** | | | | |  |  |  |  |
| **emotion** | **n** | **estimate** | **ratio** | **p.adj** |  |  |  |  |
| Anger | 647 | 0.113 | 1.120 | **0.0002** |  |  |  |  |
| Disgust | 647 | -0.016 | 0.984 | 0.061 |  |  |  |  |
| Fear | 647 | 0.090 | 1.094 | **0.002** |  |  |  |  |
| Happiness | 647 | -0.372 | 0.689 | **< 0.0001** |  |  |  |  |
| Sadness | 647 | -0.108 | 0.898 | **0.008** |  |  |  |  |
| Surprise | 647 | -0.033 | 0.968 | 0.35 |  |  |  |  |

**Supplementary Table 1. Comparison of the emotional and hedonic properties between the olfactory and non-olfactory memory.**
