## Supplementary Table 2 for "The formation and content of odor memory from childhood"

| **Sensory modalities** | |  | **Olfactory** | | **Non-Olfactory** | |
| --- | --- | --- | --- | --- | --- | --- |
| **group1** | **group2** | **df** | **statistic** | **p.adj** | **statistic** | **p.adj** |
| Audition | Gustation | 646 | 0.648 | 1 | 21.014 | **< 0.0001** |
| Audition | Olfaction | 646 | -33.030 | **< 0.0001** | 18.547 | **< 0.0001** |
| Audition | Touch | 646 | -2.031 | 0.427 | 5.151 | **< 0.0001** |
| Audition | Vision | 646 | -22.588 | **< 0.0001** | -20.010 | **< 0.0001** |
| Gustation | Olfaction | 646 | -28.884 | **< 0.0001** | -4.864 | **< 0.0001** |
| Gustation | Touch | 646 | -2.415 | 0.16 | -17.520 | **< 0.0001** |
| Gustation | Vision | 646 | -18.689 | **< 0.0001** | -45.882 | **< 0.0001** |
| Olfaction | Touch | 646 | 31.519 | **< 0.0001** | -13.978 | **< 0.0001** |
| Olfaction | Vision | 646 | 13.968 | **< 0.0001** | -41.818 | **< 0.0001** |
| Touch | Vision | 646 | -19.419 | **< 0.0001** | -24.789 | **< 0.0001** |
| **Olfactory vs Non-olfactory memory** | | | |  |  |  |
| **Sense** | **statistic** | **df** | **p.adj** |  |  |  |
| Audition | 9.310 | 646 | **< 0.0001** |  |  |  |
| Gustation | -11.892 | 646 | **< 0.0001** |  |  |  |
| Olfaction | -47.017 | 646 | **< 0.0001** |  |  |  |
| Touch | 1.579 | 646 | 0.115 |  |  |  |
| Vision | 12.010 | 646 | **< 0.0001** |  |  |  |

**Supplementary Table 2. Comparison of the relative contribution of each sensory modality to the olfactory and non-olfactory memory.**
