## Supplementary Table 3 for "The formation and content of odor memory from childhood"

| **Olfactory vs Non olfactory memory – Repetition of the initial event** | | | | | |
| --- | --- | --- | --- | --- | --- |
| **Repetition** | **N_Olf** | **N_Non_Olf** | **Prop_Olf** | **Prop_Non_Olf** | **p.adj** |
| 1 | 87 | 323 | 13.4 | 49.9 | **< 0.0001** |
| 2-5 | 87 | 87 | 13.4 | 13.4 | 1 |
| >5 | 473 | 237 | 73.1 | 36.6 | **< 0.0001** |

**Supplementary Table 3. Comparison of the frequency of repetition of the initial event between the olfactory and non-olfactory memory.** Data for the olfactory memory were already published in Dejou et al. (2026) and reused here for the purpose of comparison with the non-olfactory memory.
