## Supplementary Table 4 for "The formation and content of odor memory from childhood"

| **Olfactory memory – Odor classification** | | | | |
| --- | --- | --- | --- | --- |
| **group1** | **group2** | **n1** | **n2** | **p.adj** |
| Food | Rain & atmosphere | 190 | 31 | **< 0.0001** |
| Animal & human body odors | Food | 41 | 190 | **< 0.0001** |
| Rain & atmosphere | Nature & vegetation | 31 | 169 | **< 0.0001** |
| Food | Perfume & personal care | 190 | 42 | **< 0.0001** |
| Animal & human body odors | Nature & vegetation | 41 | 169 | **< 0.0001** |
| Perfume & personal care | Nature & vegetation | 42 | 169 | **< 0.0001** |
| Food | Industry & chemistry | 190 | 54 | **< 0.0001** |
| Industry & chemistry | Nature & vegetation | 54 | 169 | **< 0.0001** |
| Food | Home | 190 | 87 | **< 0.0001** |
| Home | Rain & atmosphere | 87 | 31 | **< 0.0001** |
| Home | Nature & vegetation | 87 | 169 | **< 0.0001** |
| Animal & human body odors | Home | 41 | 87 | **< 0.0001** |
| Home | Perfume & personal care | 87 | 42 | **< 0.0001** |
| Home | Industry & chemistry | 87 | 54 | **0.0002** |
| Industry & chemistry | Rain & atmosphere | 54 | 31 | **0.001** |
| Animal & human body odors | Industry & chemistry | 41 | 54 | 0.11 |
| Perfume & personal care | Rain & atmosphere | 42 | 31 | 0.12 |
| Industry & chemistry | Perfume & personal care | 54 | 42 | 0.13 |
| Animal & human body odors | Rain & atmosphere | 41 | 31 | 0.14 |
| Food | Nature & vegetation | 190 | 169 | 0.14 |
| Animal & human body odors | Perfume & personal care | 41 | 42 | 1.00 |

**Supplementary Table 4. Comparison of the frequency of the different odor categories in the olfactory memory.**
