## Supplementary Table 5 for "The formation and content of odor memory from childhood"

| **Olfactory memory – Who (1)** | | | | | **Olfactory memory – Who (2)** | | | | |
| --- | --- | --- | --- | --- | --- | --- | --- | --- | --- |
| **group1** | **group2** | **n1** | **n2** | **p.adj** | **group1** | **group2** | **n1** | **n2** | **p.adj** |
| Mother | Sister | 235 | 6 | **< 0.0001** | Grandfather | Childminder | 42 | 9 | **< 0.0034** |
| Mother | Childminder | 235 | 9 | **< 0.0002** | Friends | Childminder | 42 | 9 | **< 0.0035** |
| Grandmother | Mother | 11 | 235 | **< 0.0003** | Cousins | Childminder | 41 | 9 | **< 0.0036** |
| Mother | Aunt | 235 | 31 | **< 0.0004** | Grandfather | Grandmother | 42 | 11 | **< 0.0037** |
| Mother | Uncle | 235 | 29 | **< 0.0005** | Grandmother | Friends | 11 | 42 | **< 0.0038** |
| Mother | Cousins | 235 | 41 | **< 0.0006** | Grandmother | Cousins | 11 | 41 | **< 0.0039** |
| Mother | Teachers | 235 | 38 | **< 0.0007** | Childminder | Teachers | 9 | 38 | **< 0.0040** |
| Grandfather | Mother | 42 | 235 | **< 0.0008** | Sister | Aunt | 6 | 31 | **0.0001** |
| Mother | Friends | 235 | 42 | **< 0.0009** | Grandmother | Teachers | 11 | 38 | **0.0003** |
| Father | Sister | 126 | 6 | **< 0.0010** | Sister | Uncle | 6 | 29 | **0.0003** |
| Father | Childminder | 126 | 9 | **< 0.0011** | Aunt | Childminder | 31 | 9 | **0.001** |
| Grandmother | Father | 11 | 126 | **< 0.0012** | Uncle | Childminder | 29 | 9 | **0.003** |
| Brother | Sister | 97 | 6 | **< 0.0013** | Grandmother | Aunt | 11 | 31 | **0.004** |
| Brother | Childminder | 97 | 9 | **< 0.0014** | Grandmother | Uncle | 11 | 29 | **0.008** |
| Mother | Brother | 235 | 97 | **< 0.0015** | Father | Brother | 126 | 97 | 0.05 |
| Grandmother | Brother | 11 | 97 | **< 0.0016** | Uncle | Cousins | 29 | 41 | 0.13 |
| Father | Uncle | 126 | 29 | **< 0.0017** | Grandfather | Uncle | 42 | 29 | 0.20 |
| Father | Aunt | 126 | 31 | **< 0.0018** | Uncle | Friends | 29 | 42 | 0.20 |
| Father | Teachers | 126 | 38 | **< 0.0019** | Aunt | Cousins | 31 | 41 | 0.22 |
| Father | Mother | 126 | 235 | **< 0.0020** | Grandfather | Aunt | 42 | 31 | 0.29 |
| Father | Cousins | 126 | 41 | **< 0.0021** | Aunt | Friends | 31 | 42 | 0.30 |
| Grandfather | Father | 42 | 126 | **< 0.0022** | Grandmother | Sister | 11 | 6 | 0.39 |
| Father | Friends | 126 | 42 | **< 0.0023** | Uncle | Teachers | 29 | 38 | 0.39 |
| Brother | Uncle | 97 | 29 | **< 0.0024** | Aunt | Teachers | 31 | 38 | 0.54 |
| Brother | Aunt | 97 | 31 | **< 0.0025** | Sister | Childminder | 6 | 9 | 0.69 |
| Grandfather | Sister | 42 | 6 | **< 0.0026** | Friends | Teachers | 42 | 38 | 0.79 |
| Sister | Friends | 6 | 42 | **< 0.0027** | Grandfather | Teachers | 42 | 38 | 0.81 |
| Brother | Teachers | 97 | 38 | **< 0.0028** | Grandmother | Childminder | 11 | 9 | 0.88 |
| Brother | Cousins | 97 | 41 | **< 0.0029** | Cousins | Teachers | 41 | 38 | 0.88 |
| Sister | Cousins | 6 | 41 | **< 0.0030** | Uncle | Aunt | 29 | 31 | 0.88 |
| Grandfather | Brother | 42 | 97 | **< 0.0031** | Grandfather | Cousins | 42 | 41 | 1.00 |
| Brother | Friends | 97 | 42 | **< 0.0032** | Grandfather | Friends | 42 | 42 | 1.00 |
| Sister | Teachers | 6 | 38 | **< 0.0033** | Cousins | Friends | 41 | 42 | 1.00 |
| **Non olfactory vs olfactory memory - Who** | | | | | **Non olfactory vs olfactory memory - Who** | | | | |
| **Category** | **p_O** | **p_NO** | **ratio** | **p.adj** | **Category** | **p_O** | **p_NO** | **ratio** | **p.adj** |
| Grandmother | 1.74 | 0.32 | 5.50 | 0.06 | Uncle | 4.60 | 4.60 | 1.00 | 1.00 |
| Mother | 37.24 | 27.26 | 1.37 | **0.0005** | Cousins | 6.50 | 8.24 | 0.79 | 0.46 |
| Grandfather | 6.66 | 5.55 | 1.20 | 0.56 | Teachers | 6.02 | 7.92 | 0.76 | 0.46 |
| Father | 19.97 | 17.43 | 1.15 | 0.46 | Sister | 0.95 | 1.74 | 0.55 | 0.50 |
| Brother | 15.37 | 14.58 | 1.05 | 0.85 | Childminder | 1.43 | 2.85 | 0.50 | 0.29 |
| Aunt | 4.91 | 4.91 | 1.00 | 1.00 | Friends | 6.66 | 13.95 | 0.48 | **0.0002** |

**Supplementary Table 5. Comparison of the frequency of the different social categories between the olfactory and non-olfactory memory.**
