## Supplementary Table 6 for "The formation and content of odor memory from childhood"

| **Olfactory memory – Where** | | | | |
| --- | --- | --- | --- | --- |
| **group1** | **group2** | **n1** | **n2** | **p.adj** |
| Hospital | House | 7 | 384 | **< 0.0007** |
| Hospital | Nature/Environment | 7 | 293 | **< 0.0006** |
| House | School | 384 | 112 | **< 0.0005** |
| Hospital | Places | 7 | 213 | **< 0.0004** |
| Nature/Environment | School | 293 | 112 | **< 0.0003** |
| Hospital | School | 7 | 112 | **< 0.0002** |
| House | Places | 384 | 213 | **< 0.0001** |
| School | Places | 112 | 213 | **< 0.0000** |
| Nature/Environment | Places | 293 | 213 | **< 0.0001** |
| House | Nature/Environment | 384 | 293 | **< 0.0001** |
| **Non olfactory vs olfactory memory – Where** | | | | |
| **Category** | **p_O** | **p_NO** | **ratio** | **p.adj** |
| Nature/Environment | 46.43 | 34.87 | 1.33 | **< 0.0001** |
| Places | 33.76 | 26.78 | 1.26 | **0.01** |
| House | 60.86 | 52.14 | 1.17 | **0.004** |
| School | 17.75 | 23.14 | 0.77 | **0.02** |
| Hospital | 1.11 | 2.85 | 0.39 | **0.04** |

**Supplementary Table 6. Comparison of the frequency of the different location categories between the olfactory and non-olfactory memory.**
