## Supplementary Table 7 for "The formation and content of odor memory from childhood"

| **Olfactory memory – Emotions (1)** | | | | | **Olfactory memory – Emotions (2)** | | | | |
| --- | --- | --- | --- | --- | --- | --- | --- | --- | --- |
| **group1** | **group2** | **n1** | **n2** | **p.adj** | **group1** | **group2** | **n1** | **n2** | **p.adj** |
| Joy | Shame | 256 | 3 | **< 0.0001** | Surprise | Discomfort | 6 | 32 | **< 0.0040** |
| Joy | Anger | 256 | 4 | **< 0.0002** | Surprise | Fear | 6 | 32 | **< 0.0041** |
| Joy | Curiosity | 256 | 8 | **< 0.0003** | Affection | Safety | 22 | 56 | **0.0001** |
| Joy | Surprise | 256 | 6 | **< 0.0004** | Nostalgia | Curiosity | 34 | 8 | **0.0002** |
| Joy | Sadness | 256 | 17 | **< 0.0005** | Curiosity | Fear | 8 | 32 | **0.0002** |
| Joy | Affection | 256 | 22 | **< 0.0006** | Curiosity | Discomfort | 8 | 32 | **0.0003** |
| Wellbeing | Shame | 217 | 3 | **< 0.0007** | Affection | Excitement | 22 | 54 | **0.0005** |
| Wellbeing | Surprise | 217 | 6 | **< 0.0008** | Affection | Shame | 22 | 3 | **0.0005** |
| Wellbeing | Anger | 217 | 4 | **< 0.0009** | Affection | Anger | 22 | 4 | **0.001** |
| Wellbeing | Curiosity | 217 | 8 | **< 0.0010** | Sadness | Shame | 17 | 3 | **0.005** |
| Joy | Nostalgia | 256 | 34 | **< 0.0011** | Affection | Surprise | 22 | 6 | **0.007** |
| Joy | Fear | 256 | 32 | **< 0.0012** | Anger | Sadness | 4 | 17 | **0.009** |
| Wellbeing | Sadness | 217 | 17 | **< 0.0013** | Safety | Discomfort | 56 | 32 | **0.02** |
| Joy | Discomfort | 256 | 32 | **< 0.0014** | Excitement | Fear | 54 | 32 | **0.02** |
| Joy | Excitement | 256 | 54 | **< 0.0015** | Affection | Curiosity | 22 | 8 | **0.02** |
| Wellbeing | Affection | 217 | 22 | **< 0.0016** | Safety | Fear | 56 | 32 | **0.02** |
| Wellbeing | Nostalgia | 217 | 34 | **< 0.0017** | Excitement | Discomfort | 54 | 32 | **0.03** |
| Wellbeing | Fear | 217 | 32 | **< 0.0018** | Safety | Nostalgia | 56 | 34 | **0.03** |
| Joy | Safety | 256 | 56 | **< 0.0019** | Joy | Wellbeing | 256 | 217 | **0.03** |
| Wellbeing | Discomfort | 217 | 32 | **< 0.0020** | Nostalgia | Sadness | 34 | 17 | **0.03** |
| Wellbeing | Safety | 217 | 56 | **< 0.0021** | Surprise | Sadness | 6 | 17 | **0.04** |
| Wellbeing | Excitement | 217 | 54 | **< 0.0022** | Excitement | Nostalgia | 54 | 34 | **0.05** |
| Safety | Shame | 56 | 3 | **< 0.0023** | Discomfort | Sadness | 32 | 17 | 0.05 |
| Excitement | Shame | 54 | 3 | **< 0.0024** | Sadness | Fear | 17 | 32 | 0.05 |
| Safety | Anger | 56 | 4 | **< 0.0025** | Curiosity | Sadness | 8 | 17 | 0.13 |
| Excitement | Anger | 54 | 4 | **< 0.0026** | Affection | Nostalgia | 22 | 34 | 0.16 |
| Safety | Surprise | 56 | 6 | **< 0.0027** | Affection | Discomfort | 22 | 32 | 0.25 |
| Excitement | Surprise | 54 | 6 | **< 0.0028** | Affection | Fear | 22 | 32 | 0.26 |
| Excitement | Curiosity | 54 | 8 | **< 0.0029** | Curiosity | Shame | 8 | 3 | 0.26 |
| Safety | Curiosity | 56 | 8 | **< 0.0030** | Curiosity | Anger | 8 | 4 | 0.44 |
| Nostalgia | Shame | 34 | 3 | **< 0.0031** | Surprise | Shame | 6 | 3 | 0.56 |
| Discomfort | Shame | 32 | 3 | **< 0.0032** | Affection | Sadness | 22 | 17 | 0.57 |
| Fear | Shame | 32 | 3 | **< 0.0033** | Surprise | Anger | 6 | 4 | 0.81 |
| Nostalgia | Anger | 34 | 4 | **< 0.0034** | Curiosity | Surprise | 8 | 6 | 0.84 |
| Anger | Fear | 4 | 32 | **< 0.0035** | Nostalgia | Discomfort | 34 | 32 | 0.94 |
| Discomfort | Anger | 32 | 4 | **< 0.0036** | Nostalgia | Fear | 34 | 32 | 0.94 |
| Safety | Sadness | 56 | 17 | **< 0.0037** | Excitement | Safety | 54 | 56 | 0.94 |
| Excitement | Sadness | 54 | 17 | **< 0.0038** | Discomfort | Fear | 32 | 32 | 1.00 |
| Nostalgia | Surprise | 34 | 6 | **< 0.0039** | Anger | Shame | 4 | 3 | 1.00 |
| **Non olfactory vs olfactory memory - Emotions** | | | | | **Non olfactory vs olfactory memory - Emotions** | | | | |
| **Category** | **p_O** | **p_NO** | **ratio** | **p.adj** | **Category** | **p_O** | **p_NO** | **ratio** | **p.adj** |
| Safety | 8.87 | 3.33 | 2.67 | **< 0.0001** | Curiosity | 1.27 | 2.06 | 0.62 | 0.39 |
| Nostalgia | 5.39 | 2.69 | 2.00 | **0.03** | Anger | 0.63 | 2.22 | 0.29 | 0.06 |
| Wellbeing | 34.39 | 18.70 | 1.84 | **< 0.0001** | Sadness | 2.69 | 9.51 | 0.28 | **< 0.0001** |
| Affection | 3.49 | 2.38 | 1.47 | 0.38 | Fear | 5.07 | 17.91 | 0.28 | **< 0.0001** |
| Joy | 40.57 | 32.01 | 1.27 | **0.0022** | Shame | 0.48 | 1.74 | 0.27 | 0.09 |
| Excitement | 8.56 | 7.92 | 1.08 | 0.75 | Surprise | 0.95 | 3.65 | 0.26 | **0.004** |
| Discomfort | 5.07 | 6.81 | 0.74 | 0.29 |  |  |  |  |  |

**Supplementary Table 7. Comparison of the frequency of the different emotional categories between the olfactory and non-olfactory memory.**
