## Supplementary Table 8 for "The formation and content of odor memory from childhood"

|  | **Overall** | | **Less than yearly** | | **Yearly** | | **Monthly** | | **Weekly** | |
| --- | --- | --- | --- | --- | --- | --- | --- | --- | --- | --- |
|  | **X2** | **p.adj** | **X2** | **p.adj** | **X2** | **p.adj** | **X2** | **p.adj** | **X2** | **p.adj** |
| Never  vs Sometimes | 96.24 | **< 0.0001** | 23.61 | **< 0.0001** | 38.37 | **< 0.0001** | 24.24 | **< 0.0001** | 6.10 | 0.0813 |
| Never  vs Often | 125.45 | **< 0.0001** | 26.79 | **< 0.0001** | 49.50 | **< 0.0001** | 40.05 | **< 0.0001** | 6.10 | 0.0813 |
| Never  vs Always | 404.79 | **< 0.0001** | 146.69 | **< 0.0001** | 201.98 | **< 0.0001** | 52.90 | **< 0.0001** | 4.90 | 0.1611 |
| Sometimes  vs Often | 2.22 | 0.8193 | 0.03 | 1 | 0.69 | 1 | 2.03 | 0.9243 | 0.00 | 1 |
| Sometimes  vs Always | 143.29 | **< 0.0001** | 66.50 | **< 0.0001** | 91.85 | **< 0.0001** | 6.43 | 0.0674 | 0.00 | 1 |
| Often  vs Always | 110.35 | **< 0.0001** | 61.59 | **< 0.0001** | 76.01 | **< 0.0001** | 0.91 | 1 | 0.00 | 1 |

**Supplementary Table 8. Comparison of the frequency of odor-evoked memory depending on the frequency of odor exposure for the olfactory memory.**
