## Supplementary Table 9 for "The formation and content of odor memory from childhood"

| **Olfactory memory – Age effect (emotion)** | | | |
| --- | --- | --- | --- |
| **variable** | **group1** | **group2** | **p.adj** |
| Anger | 18-60 | 61-90 | **< 0.0001** |
| Disgust | 18-60 | 61-90 | **0.0006** |
| Fear | 18-60 | 61-90 | **0.005** |
| Happiness | 18-60 | 61-90 | 0.07 |
| Pleasantness | 18-60 | 61-90 | **0.04** |
| Sadness | 18-60 | 61-90 | 0.12 |
| Surprise | 18-60 | 61-90 | **< 0.0001** |

**Supplementary Table 9. Comparison of the emotional ratings of the olfactory memory between 18-60 and 61-90 age groups.**
