## Supplementary Table 10 for "The formation and content of odor memory from childhood"

|  | **Olfactory memory** | | **Non-olfactory memory** | |
| --- | --- | --- | --- | --- |
|  | Indirect | Direct | Indirect | Direct |
| Olfaction | **0.0004** | **.035** | **0.0004** | 0.075 |
| Gustation | **< 0.0001** | .80 | **< 0.0001** | **< 0.0001** |
| Vision | **0.0016** | .93 | **< 0.0001** | 0.75 |
| Audition | **0.003** | .92 | **< 0.0001** | **0.015 (neg)** |
| Touch | **< 0.0001** | **.093 (neg)** | **< 0.0001** | 0.642 |

**Supplementary Table 10. Mediation analyses of the effects of the different sensory modalities on the pleasantness of olfactory and non-olfactory memories.**
